# Sleep modulates alcohol toxicity in *Drosophila*

**DOI:** 10.1101/2021.04.16.440198

**Authors:** Eric J. Noakes, Aliza K. De Nobrega, Alana P. Mellers, Lisa C. Lyons

**Author notes:** Corresponding Author with complete address: Lisa C. Lyons Department of Biological Science Florida State University 319 Stadium Drive Tallahassee, FL 32306-4295. These authors contributed equally (Eric J. Noakes, Aliza K. De Nobrega).

## Abstract

**Study Objectives:** Alcohol abuse is a significant public health problem, particularly in populations in which sleep deprivation is common as such as shift workers and aged individuals. Although research demonstrates the effect of alcohol on sleep, little is known about the role of sleep in alcohol sensitivity and toxicity. We investigated sleep as a factor modulating alcohol toxicity using *Drosophila melanogaster*, a model system ideal for studies of sleep, alcohol and aging.

**Methods:** Following 24 hours of sleep deprivation using mechanical stimulation, *Drosophila* were exposed to binge-like alcohol exposures. Behavioral sensitivity, tolerance, and mortality were assessed. The effects of chronic sleep deprivation on alcohol toxicity were investigated using a short sleep mutant *insomniac*. Pharmacological induction of sleep for prior to alcohol exposure was accomplished using a GABA_A_-receptor agonist, 4,5,6,7-tetrahydroisoxazolo(5,4-c)pyridin-3-ol (THIP) to determine if increased sleep mitigated the effects of alcohol toxicity on middle-aged flies and flies with environmentally disrupted circadian clocks mimicking groups more vulnerable to the effects of alcohol.

**Results:** Acute sleep deprivation increased alcohol-induced mortality following alcohol exposure. However, sleep deprivation had no effect on alcohol absorbance or clearance. Sleep deprivation also abolished functional tolerance measured 24 hours after the initial alcohol exposure, although tolerance at 4 h was observed. Pharmacologically increasing sleep prior to alcohol exposure decreased alcohol-induced mortality.

**Conclusions:** Sleep quantity prior to alcohol exposure affects alcohol toxicity with decreased sleep increasing alcohol toxicity and dampened 24-hour alcohol tolerance. In contrast, increased sleep mitigated alcohol-induced mortality even in vulnerable groups such as aging flies and those with circadian dysfunction.

**Statement of significance:** With the growing incidence of sleep deprivation and sleep disorders across adolescents and adults, it is important to understand the role of sleep in alcohol toxicity to develop future therapies for prevention and treatment of alcohol-induced pathologies. Using *Drosophila melanogaster*, an established model for both sleep and alcohol research, we found that acute and chronic sleep deprivation increased alcohol toxicity and eliminated long-term functional alcohol tolerance. In contrast, increased sleep prior to binge-like alcohol exposure mitigated alcohol-induced mortality even in vulnerable groups with higher susceptibility to alcohol toxicity.

## Introduction

Alcohol abuse and its associated pathologies is a pervasive societal problem with serious negative impacts on individual health, family structure and the economy^1–8^. In the United States alcohol use disorders account for 79% of all diagnoses of substance use disorders^9^ and the economic impact of alcohol misuse is estimated at $249 billion annually^7, 10^. Alcohol abuse and alcohol pathologies appear higher in populations in which sleep deprivation is common including teenagers, young adults, shift workers and aged individuals^11–18^. Although considerable behavioral research has demonstrated the effects of alcohol on sleep homeostasis^19–21^, surprisingly little is known about the role of sleep in modulating alcohol sensitivity and toxicity at the physiological level. Sleep impairments are traditionally viewed as symptoms of alcohol use disorders; however, sleep disorders increase the incidence and risk of relapse in recovering alcoholics^22–25^. Sleep deprivation represents a significant rising public health problem in the United States and the world^26–29^. The pervasiveness of factors contributing to sleep disruptions including artificial light at night, the use of personal electronics and the increase in shiftwork and extended work days^30, 31^, combined with the increased risk of substance abuse associated with sleep deprivation, makes understanding how sleep deprivation affects alcohol-induced behaviors and toxicity imperative to identify and optimize therapies for future prevention and treatment of alcohol-induced pathologies.

The high degree of physiological, molecular and neurological conservation between the fruit fly *Drosophila melanogaster* and mammals makes *Drosophila* an ideal model for the investigation of sleep and alcohol interactions^32–34^. Sleep in *Drosophila* occurs in stages, varying in intensity during the night with observable sex and age dependent differences^35–44^. As in other species, circadian and homeostatic processes regulate sleep in flies with waking activity affecting sleep need^38, 45^. Moreover, alcohol physiology is remarkably conserved from flies to humans with parallels in behaviors as well as the underlying molecular mechanisms^46, 47^. When exposed to alcohol vapor, initially flies exhibit hyperactivity with increased locomotor activity, followed by a loss of motor control and eventually sedation^48–51^. Flies also develop functional alcohol tolerance dependent upon changes in neural plasticity^50, 52–55^ and addiction-like behaviors^56–58^ with a preference for alcohol following previous exposure^59, 60^.

We investigated the role of decreased and increased sleep in modulating alcohol toxicity. We found that acute sleep deprivation increased behavioral sensitivity and mortality following acute and repeated exposure to alcohol. These effects were independent of alcohol metabolism as no differences were observed in alcohol absorption and clearance between sleep deprived and non-sleep deprived flies. Sleep deprivation also inhibited the induction of long-term functional alcohol tolerance observed 24 h following the first alcohol exposure, although short-term tolerance measured 4 h following the first alcohol exposure was not affected. Chronic sleep restriction also increased alcohol-induced mortality. Encouragingly, we found that pharmacologically increasing sleep had the opposite effect of sleep deprivation, ameliorating alcohol mortality in middle-aged flies and flies with a disrupted circadian clock. This research highlights the critical role of sleep as a factor in alcohol toxicity.

## Methods

### Fly Maintenance

All flies were maintained on standard cornmeal-molasses food at 25°C and 60-70% relative humidity in 12:12 light-dark (LD) cycles. *Insomniac* (*inc*) mutants and the background *w^1118^* line were generously provided by Dr. Nicholas Stavropoulos, New York University. Adult flies (∼30 per vial) were transferred approximately every 3 days to maintain stress-free cultures. All experiments were carried out in an environmentally controlled dark room at 25°C and 60-70% relative humidity under dim red light. Zeitgeber time (ZT) 0 represents lights on and ZT 12 corresponds with lights off. For experiments performed in constant light (LL) conditions, flies were transferred to LL on the day of eclosion.

### Alcohol Exposure

Alcohol vapor exposure was performed as previously described^51, 61, 62^. Four tubes, each containing ∼30 flies, received a steady flow of ethanol vapor at a pre-determined percentage. Precise alcohol percentages were achieved by mixing air bubbled through deionized water and 95% ethanol (Koptec, Declon Labs, Inc.). Air flow rates were monitored throughout the experiment to ensure consistency of alcohol concentration. Water vapor controls were run simultaneously with 100% water vapor. Alcohol exposures were performed at ZT 9 to avoid circadian variation in responses unless otherwise stated for a specific protocol.

### Sleep Deprivation

Consistent sleep deprivation was achieved using gentle mechanical stimulation on the GyroMini Nutating Mixer (Labnet International, Inc.). Vials containing ∼30 flies were placed at an angled position in a larger beaker with a raised block at a fixed position inside the beaker. Mixer rotation caused the vials to rotate within the beaker and then gently jump over the raised block, providing the flies with a startle movement every 2.5 seconds. The constant motion of the vials combined with the startle ensured consistent sleep deprivation with no apparent injuries or increased mortality observed after 24 hours of sleep deprivation. Sleep deprivation was performed in an incubator under 25°C, 60-70% relative humidity and 12:12 LD conditions. Non-sleep deprived controls were housed in the same incubator.

### Sedation

Alcohol-induced sedation was performed as previously described^48^. Briefly, flies were exposed to 50% alcohol vapor for one hour with observations of behavioral state made every five minutes following a gentle tap of the vial. Flies were scored as sedated when immobile and lacking coordinated leg movements except for spontaneous twitching^52^. The mean time to 50% sedation was calculated using a linear extrapolation.

### Tolerance

Tolerance was determined as previously described^51^. Flies received a pre-exposure of 50% alcohol for 30 minutes at ZT4.5 following a one-hour dark room acclimation period. Sedation was assessed during the pre-exposure. Flies were then returned to food vials to allow time for recovery and complete metabolism of the alcohol before testing. Testing occurred 4 hours later at ZT9 for short-term rapid tolerance or 24 hours later for long-term rapid tolerance with all experimental groups represented at each test. Tolerance was defined as an increase in average time to reach 50% sedation from the pre-exposure with the difference in sedation time between naïve and pre-exposed flies used for quantification. For tolerance experiments, sleep deprivation took place from ZT3.5 – ZT3.5.

### Mortality

Following each alcohol exposure, flies were returned to food vials placed horizontally for approximately 2 h to allow recovery of postural control. Immediate mortality was assessed 24 hours following the last alcohol exposure and then daily for 6 days. Delayed mortality refers to the cumulative mortality within seven days of the final alcohol exposure.

### Gaboxadol Treatment

Sleep was pharmacologically increased with the GABA-A agonist, 4,5,6,7-tetrahydroisoxazolo [5,4-c]pyridin-3-ol (THIP or Gaboxadol). 10 d old (in constant light) or 20 d old (in LD) flies were transferred to Gaboxadol-containing food (0.05 mg/mL) for either 48 or 24 h respectively prior to repeated alcohol exposure. A repetitive alcohol exposure protocol was used to assess alcohol-induced mortality as described previously^61^. Flies were exposed for 3 days to 1 h alcohol vapor at ZT 9 (exposures separated by 24 h).

### Alcohol Absorbance

Following 24 h sleep deprivation, batches of 20 flies were exposed to 50% alcohol vapor for 30 minutes at ZT 9, after which they were frozen at 0, 0.5, 1, 2, or 4 h following alcohol exposure. Alcohol absorbance was measured using an enzymatic alcohol dehydrogenase assay (ADH-NAD kit; Sigma-Aldrich) per the manufacturer’s directions and as described previously^49, 51^. Briefly, flies were homogenized in 200 uL refrigerated Tris-HCl (pH 7.5) buffer. Homogenate was spun at 15,000 x g for 20 min at 4°C. 250 uL NAD-ADH reagent was added to a 5-µL aliquot of supernatant. Absorbance was measured at 340 nm within 20 min using a 96-well plate format and a Versa-Max plate reader (brand). Alcohol absorbance was normalized to total protein to eliminate the effect of body size variation between batches of flies.

### Locomotor Activity Rhythms

Locomotor activity of adult male flies was monitored using *Drosophila* activity monitors (Trikinetics, Waltham, MA) as described previously^63^. Sleep activity of flies was recorded following entrainment either in LL or LD cycles at 25°C for 4 days, after which the flies were transferred without using anesthetic to Gaboxadol-containing media for 48 hours for further measurements. Sleep data were analyzed using the ClockLab Suite.

### Statistics

Statistics were performed using GraphPad Prism Version 6.0. Experimental groups were compared using analysis of variance (ANOVA). Post-hoc analyses in multiple comparisons were performed using the Bonferroni correction.

## Results

### Sleep deprivation increases sensitivity to alcohol-induced sedation

Sleep deprivation appears to be a contributing factor to the increased use of alcohol as suggested in studies of shift workers and young adults^64–69^. However, relatively little is known about the effects of sleep deprivation on alcohol toxicity and alcohol pathologies. As a first step in exploring the modulatory role of sleep on alcohol sensitivity, we evaluated the effect of 24 hours sleep loss on alcohol sensitivity. *Drosophila* (mixed sex, 10 d old) were sleep deprived using mechanical sleep deprivation for 24 hours (ZT 8 – ZT 8) and then exposed to 50% alcohol vapor (1 h exposure; Figure 1A). As predicted given the sedative effects of alcohol, sleep deprived flies were sedated significantly faster than non-sleep deprived age-matched controls indicating that sleep loss increases behavioral sensitivity to alcohol (Figure 1B – 1C; *t*_(14)_ = 13.46, *p* = 0.0002).

**Figure 1:**
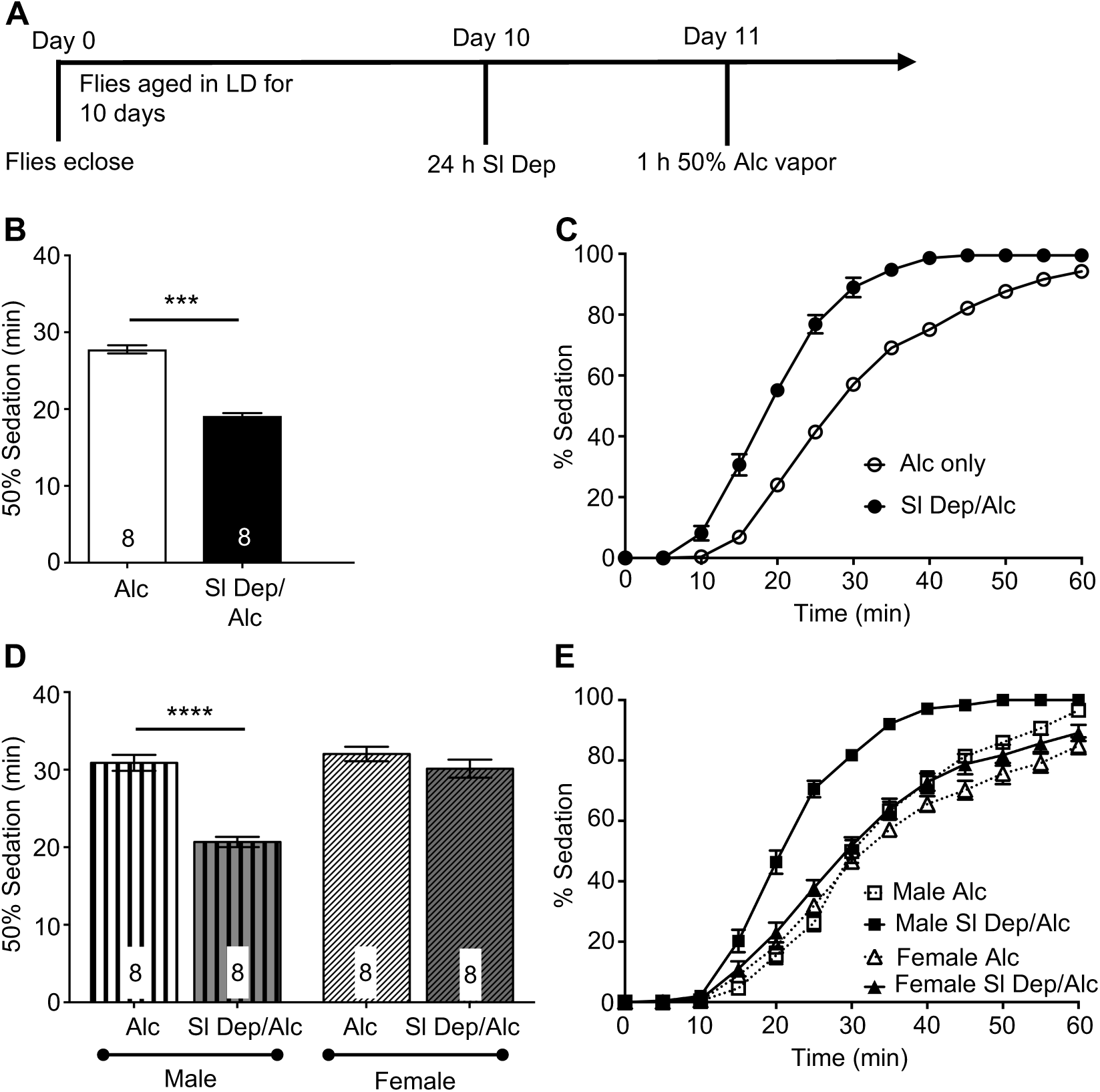
Acute sleep deprivation increases sensitivity to alcohol-induced sedation. A) 10 d old WT CS flies were sleep – deprived for 24 h then exposed to 50% alcohol vapor for 1 h. Sensitivity to sedation was measured by counting the number of flies sedated every 5 min. B) Sleep deprivation significantly exacerbates alcohol-induced sedation (*t*_(14)_ = 13.4558, *p* = 0.0002). Mean time necessary for 50% of the flies to become sedated during alcohol exposure and standard error of the mean plotted for all experiments. C) Complete time course of alcohol exposure showing percent of flies exhibiting sedation for 10 d old sleep-deprived and non-sleep deprived flies. D) Separate groups of 10 d male and female flies were sleep deprived for 24 h then exposed to 50% alcohol vapor for 1 h. Sleep-deprived males sedate faster compared to sleep-deprived females, indicating that increased sensitivity to alcohol sedation is not due to an exacerbated response to stress. (ANOVA *F*_3,28_ = 29.24, *p* < 0.0001). N shown on bars for each group is the number of vials tested for each group with 25-30 flies per vial. E) Complete time course of alcohol exposure showing percent of flies exhibiting sedation for 10 d old sleep-deprived and non-sleep deprived male and female flies.

Potentially, the alteration in alcohol sensitivity between the groups may have arisen from the mechanical procedure used for sleep deprivation rather than sleep deprivation itself. To test whether sleep deprivation or mechanically induced stress was the underlying cause of the observed difference in alcohol sensitivity, we took advantage of the sex-specific difference in daytime sleep in flies^38^. Male flies exhibit a daytime “siesta sleep” period in which they sleep significantly more during the daytime as compared to mated females^70^. If the increased behavioral alcohol sensitivity observed in sleep deprived flies was attributed to sleep loss, sleep deprivation during the daytime should have a greater impact on males. However, if the increased sensitivity to alcohol were due to mechanically induced stress, the effects of daytime sleep deprivation would be similar in males and females. Mated 10-d old female and male flies were separately sleep deprived for the first eight hours of the subjective day (ZT 0 – ZT 8) and exposed to 50% alcohol vapor for 1 h at ZT 9 with sedation assayed at 5 min intervals. Sleep deprived males were significantly more sensitive to alcohol than non-sleep deprived males with shorter exposure times inducing sedation (Figure 1D – 1E; ANOVA: *F_3,28_* = 29.24, *p* < 0.0001). In contrast, sleep deprivation appeared to have little effect on female flies with sleep deprived females showing similar alcohol responses to non-sleep deprived flies (Figure 1D – 1E; ANOVA: *F_3,28_* = 29.24, *p* < 0.0001). These results suggest that sleep deprivation increases the behavioral sensitivity to alcohol and these effects are independent of any stress from mechanical perturbation.

### Sleep deprivation increases acute and chronic alcohol toxicity

Excessive binge drinking escalates the incidence of alcohol-poisoning deaths^71, 72^. Therefore, it is important to understand the potential confounding effects of sleep loss on alcohol toxicity. To determine whether sleep deprivation alters alcohol toxicity, we tested the effect of a single exposure to 50% alcohol vapor on mortality. Flies (10 d, mixed sex) were sleep deprived for 24 h (ZT 8 – ZT 8) and exposed to 50% alcohol vapor for one h at ZT 9 (Figure 2A). Nearly 100% of the flies become sedated under this protocol. Mortality was assessed 24 h and 7 days following alcohol exposure. When exposed to alcohol vapor following sleep deprivation, flies showed significant mortality 24 h after alcohol exposure compared to non-sleep deprived flies exposed to alcohol vapor or flies that were sleep deprived and exposed to water vapor (Figure 2B, ANOVA: *F_3,28_* = 22.50, *p* < 0.0001 and 2C; ANOVA: *F_3,28_* = 14.01, *p* < 0.0001). However, there was no significant further increase in alcohol induced mortality when cumulative delayed mortality was measured 7 days following the exposure to alcohol suggesting that the effects of sleep deprivation on alcohol toxicity occurred within the first 24 hours. Non-sleep deprived and sleep deprived flies exposed to water vapor alone had negligible levels of mortality at either 24 h or 7 days suggesting that 24 h sleep deprivation itself does not result in mortal injury to the flies (Figure 2B and 2C). Exposure to alcohol vapor caused a noticeable but not significant rise in immediate and delayed mortality compared to water vapor controls (Figure 2B and C). These results suggest that sleep deprivation exacerbates the acute toxicity of alcohol with primary mortality observed within 24 hours of alcohol exposure (Figure 2B). We next investigated the effects sleep deprivation prior to a repeat binge alcohol exposure paradigm. As previously, flies were sleep deprived for 24 h (ZT 8 – ZT 8) and then exposed to 40% alcohol vapor for 1 h (ZT 9) on 3 consecutive days (Figure 2D). Perhaps not surprisingly, the first alcohol exposure after sleep deprivation induced a significant increase in mortality (Figure 2E, ANOVA: *F_3,76_* = 15.42, *p* < 0.0001). Alcohol-induced mortality was not significantly higher following the 2^nd^ and 3^rd^ alcohol exposures (Figure 2F), potentially due to the opportunity for recovery sleep following the 1^st^ exposure to alcohol. The degree of mortality observed 7 days following the last alcohol exposure was similar between the acute and repeated binge alcohol paradigms (Figure 2G, ANOVA: *F_3,76_* = 19.91, *p* < 0.0001).

**Figure 2:**
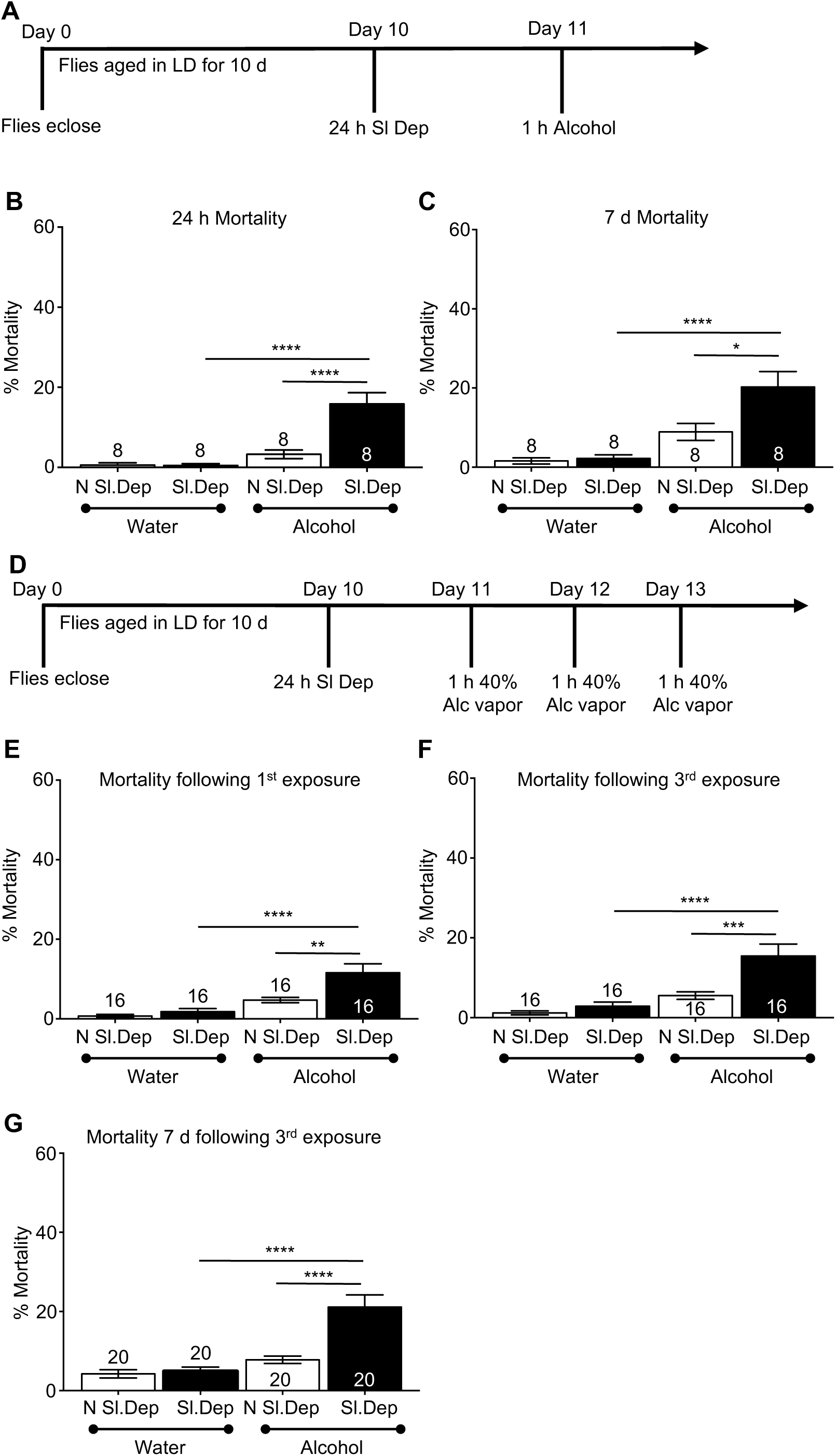
Acute sleep deprivation exacerbates alcohol-induced mortality following a single and repeated exposures to alcohol. A) 10 d old WT CS flies were sleep – deprived for 24 h then exposed to 50% alcohol vapor for 1 h. The number of flies that died were counted every 24 h for 7 days following alcohol exposure. B) Sleep-deprived flies exhibited a drastic increase in mortality within 24 h of exposure to alcohol compared to non-sleep deprived flies (ANOVA *F*_3,28_ = 22.50, *p* < 0.0001). C) Mortality continued to significantly rise 7 days following exposure to alcohol in both non-sleep deprived and sleep deprived flies (ANOVA: *F_3,28_* = 14.01, *p* < 0.0001). D) 10 d old WT CS flies were sleep – deprived for 24 h followed by 3 consecutive exposures to 1 h alcohol (50% alcohol vapor) at ZT 9 with each exposure separated by 24 h. E) Sleep-deprived flies exhibited a drastic increase in mortality within 24 h of 1^st^ exposure to alcohol compared to non-sleep deprived flies (ANOVA: *F_3,76_* = 15.42, *p* < 0.0001). F) Mortality continued to rise 24 h following the 3^rd^ exposure to alcohol vapor in both non-sleep deprived and sleep deprived flies (ANOVA: *F_3,76_* = 14.42, *p* < 0.0001), although this rise was not significant. G) Mortality measured 7 d following the 3^rd^ alcohol exposure was significantly higher than mortality following the 1^st^ alcohol exposure (ANOVA: *F_3,76_* = 19.91, *p* < 0.0001).

### Sleep deprivation does not affect the rate of alcohol clearance

It is possible that the increases in alcohol sensitivity and mortality observed following sleep deprivation were due to increased alcohol absorption or a decline in the rate of alcohol clearance resulting in greater alcohol exposure and subsequent toxicity. To investigate this possibility, flies were sleep deprived as previously described and exposed to 50% alcohol vapor for 30 min at ZT 9 and alcohol absorbance was measured (Figure 3A). There were no significant differences in alcohol absorbance or clearance between sleep deprived flies and non-sleep deprived flies (Figure 3B). These results suggest that potential metabolic changes due to sleep deprivation do not account for the observed increased sensitivity to alcohol with the more likely possibilities including sleep deprivation induced changes in neuroadaptation at the molecular or cellular levels.

**Figure 3:**
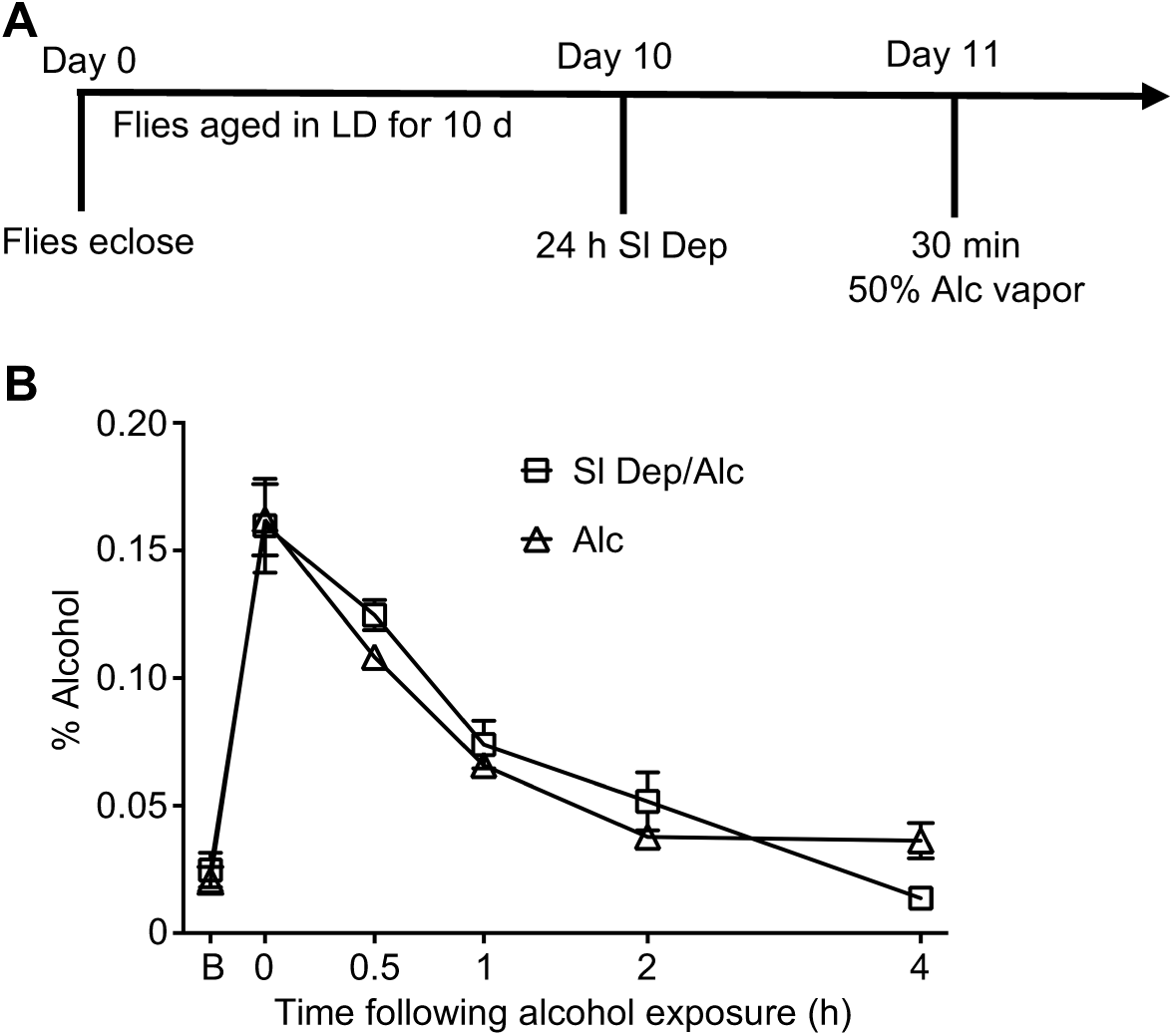
Sleep deprivation does not affect alcohol accumulation or rate of alcohol clearance. A) Flies are aged in 12:12 LD cycle for 10 days then sleep deprived for 24 h. 10 d CS flies are exposed to 30 min of 50% alcohol vapor on day 11 with alcohol absorbance and rate of alcohol clearance assessed. B) No significant differences exist in alcohol absorbance or rate of alcohol clearance between sleep deprived and non-sleep deprived flies (n=4 per group).

### Chronic sleep deprivation induces increased alcohol-induced mortality

Chronic sleep deprivation with multiple short sleep nights may also be a predisposing factor for increased alcohol consumption and other recreational drug use^18, 73^. In the United States, approximately 70 million Americans suffer from chronic sleep loss with serious consequences for health and longevity as well as economic productivity^29, 74, 75^. To investigate the effects of chronic sleep restriction on alcohol neurobiology, we used a genetic approach rather than a mechanical system to induce sleep deprivation to avoid the possibility of stress arising from long-term mechanical stimulation. Numerous mutants with short sleep phenotypes have been identified in *Drosophila*. However, the circadian clock also regulates aspects of sleep and sleep timing, and many sleep mutants have circadian phenotypes. Given previous research demonstrating circadian modulation of alcohol sensitivity and increased alcohol-induced mortality with circadian disruption^48, 61^, we used the mutant *insomniac* that has normal circadian rhythms but exhibits a short sleep phenotype^76^ to investigate the effects of chronic sleep restriction on alcohol toxicity. *Insomniac* (*inc*) is a mutation in a putative adaptor protein for the *Cullin-3* ubiquitin ligase complex^76^. We used two *inc* mutant lines, *inc^1^* and *inc^2^* (generous gifts of Nicholas Stavropoulos at NYU), to test the effects of chronic sleep restriction on alcohol sensitivity and alcohol-induced mortality. *Both inc^1^*and *inc2* mutant lines have a 90% reduction in *inc* transcript mRNA levels with no detectable protein produced^76^. Confirming previously published results, we found that that *inc^1^* and *inc^2^* flies (Figure 4A – C) exhibit considerable reductions in total sleep time with *inc^1^* flies sleeping a little over 300 minutes per day and *inc^2^* flies sleeping approximately 600 min per day (Figure 4A, ANOVA: *F_2,67_* = 81.13, *p* < 0.0001). These mutants exhibit significantly shortened sleep bouts (Figure 4B, ANOVA: *F_2,67_* = 17.52, *p* < 0.0001) reflecting a decrease in sleep consolidation, although they do have a greater number of sleep bouts (Figure 4C, ANOVA: *F_2,67_* = 8.64, *p* < 0.0001).

**Figure 4:**
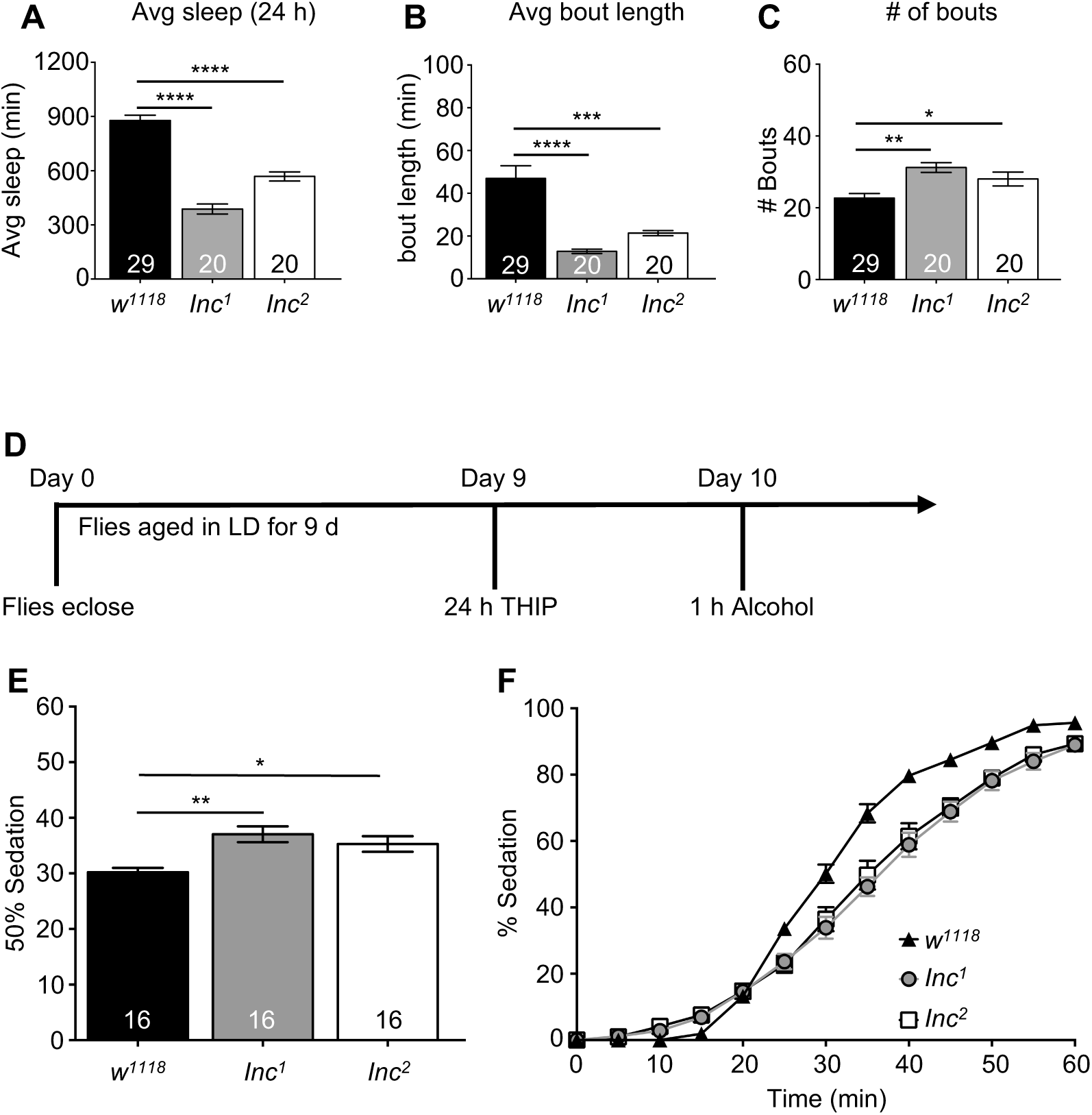
Mutations in *insomniac* do not increase behavioral sensitivity to alcohol exposure. The impact of sleep loss due to mutations in the *inc^1^* and *inc^2^* genes on alcohol sensitivity was assessed. A-C) Sleep profiles of *w^1118^*, *inc^1^* and *inc^2^* flies. *inc^1^*, *inc^2^* and flies have significantly shorter daily sleep times (A, ANOVA: *F_2,67_* = 81.13, *p* < 0.0001), shorter bout lengths (B, ANOVA: *F_2,67_* = 17.52, *p* < 0.0001) and increased number of bouts compared to *w^1118^* flies (C, ANOVA: *F_2,67_* = 8.64, *p* < 0.0001). D) 10 d *inc^1^*, *inc^2^* and *w^1118^* control flies were exposed to 50% alcohol vapor for 1 h at ZT 9 with sedation assessed every 5 minutes during the alcohol exposure. E) *inc^1^* and *inc^2^* flies demonstrated increased resistance to alcohol vapor compared to *w^1118^* controls (ANOVA: *F_2,34_* = 47.28, *p* < 0.0001). F) The complete time course for *w^1118^*, *inc^1^* and *inc^2^* flies.

To investigate the effects of chronic sleep restriction on alcohol sensitivity, we exposed 10 d old *inc^1^* and *inc^2^* flies to 50% alcohol for 1 h at ZT 9 with sedation assessed at 5-minute intervals (Figure 4D). Surprisingly, *inc^1^* and *inc^2^* flies did not exhibit increased sensitivity to alcohol; indeed, these mutants were more resistant to the sedating effects of alcohol with significantly longer times to reach 50% sedation than the *w^1118^* control flies (Figure 4D and 4E, ANOVA: *F_2,34_* = 47.28, *p* < 0.0001 with post-hoc analysis identifying significant differences between *w^1118^* vs. *inc^1^* and *w^1118^* vs *inc^2^*). These results suggest that either compensatory mechanisms exist to buffer against increased sensitivity to alcohol in these mutants or chronic sleep loss associated with the disruption of the *Cullin-3* ubiquitin ligase complex does not increase alcohol sensitivity.

While the chronic sleep deficit associated with the disruption of the *Cullin-3* ubiquitin ligase complex in the *inc* mutants did not increase alcohol sensitivity, we hypothesized that it would still increase alcohol toxicity as alcohol affects multiple signaling pathways both in the central nervous system and in peripheral tissues. To test this, we gave 10 d *inc^1^* and *inc^2^* flies a single exposure to 50% alcohol vapor for 1 h at ZT 9 and assessed mortality 24 h and 7 d following the alcohol exposure. Both *inc^1^* and *inc^2^* flies exhibited significantly higher mortality immediately (24 h) and 7 d following the exposure compared to *w^1118^* background controls (Figure 5B, ANOVA: *F_2,29_* = 16.46, *p* < 0.0001 and Figure 5C, ANOVA: *F_2,29_* = 20.87, *p* < 0.0001). Given that the *inc* mutants are postulated to have defects in ubiquitination that may affect many target proteins and signaling pathways, it is possible that the observed mortality was due to other consequences of the mutation and not to the effects of chronic sleep restriction on alcohol toxicity. Presumably as *inc* flies age, the short sleep phenotype results in an accumulated sleep debt. If this is the case, we hypothesized that younger flies (3 d) would show lower alcohol-induced mortality at levels similar to that seen with acute sleep deprivation. To test this hypothesis, we exposed 3 d *inc^1^* and *inc^2^*flies to alcohol and assessed mortality 24 h and 7 d following exposure. While there was higher mortality observed in 3 d *inc^1^* and *inc^2^* flies following alcohol exposure than 3 d *w^1118^* control flies (Figure 5E, ANOVA: *F_2,34_* = 15.03, *p* < 0.0001 and Figure 5F, ANOVA: *F_2,34_* = 25.01, *p* < 0.0001), the 3 d *inc^1^* and *inc^2^* flies exhibited significantly lower mortality following a single exposure to alcohol than 10 d *inc^1^* and *inc^2^* flies (Figure 5E - F). No significant differences were observed in alcohol-induced mortality between 3 d and 10 d *w^1118^* flies. These results suggest that the increase in alcohol-induced mortality in the 10 d *inc* mutants was due to the accrued sleep debt in the older flies rather than a non-sleep related consequence of the mutation. Together, these results suggest that separate mechanisms mediate the behavioral sensitivity to alcohol and the alcohol’s toxic effects whereby *insomniac* is necessary for the resistance to alcohol-induced mortality but not alcohol behavioral sensitivity.

**Figure 5:**
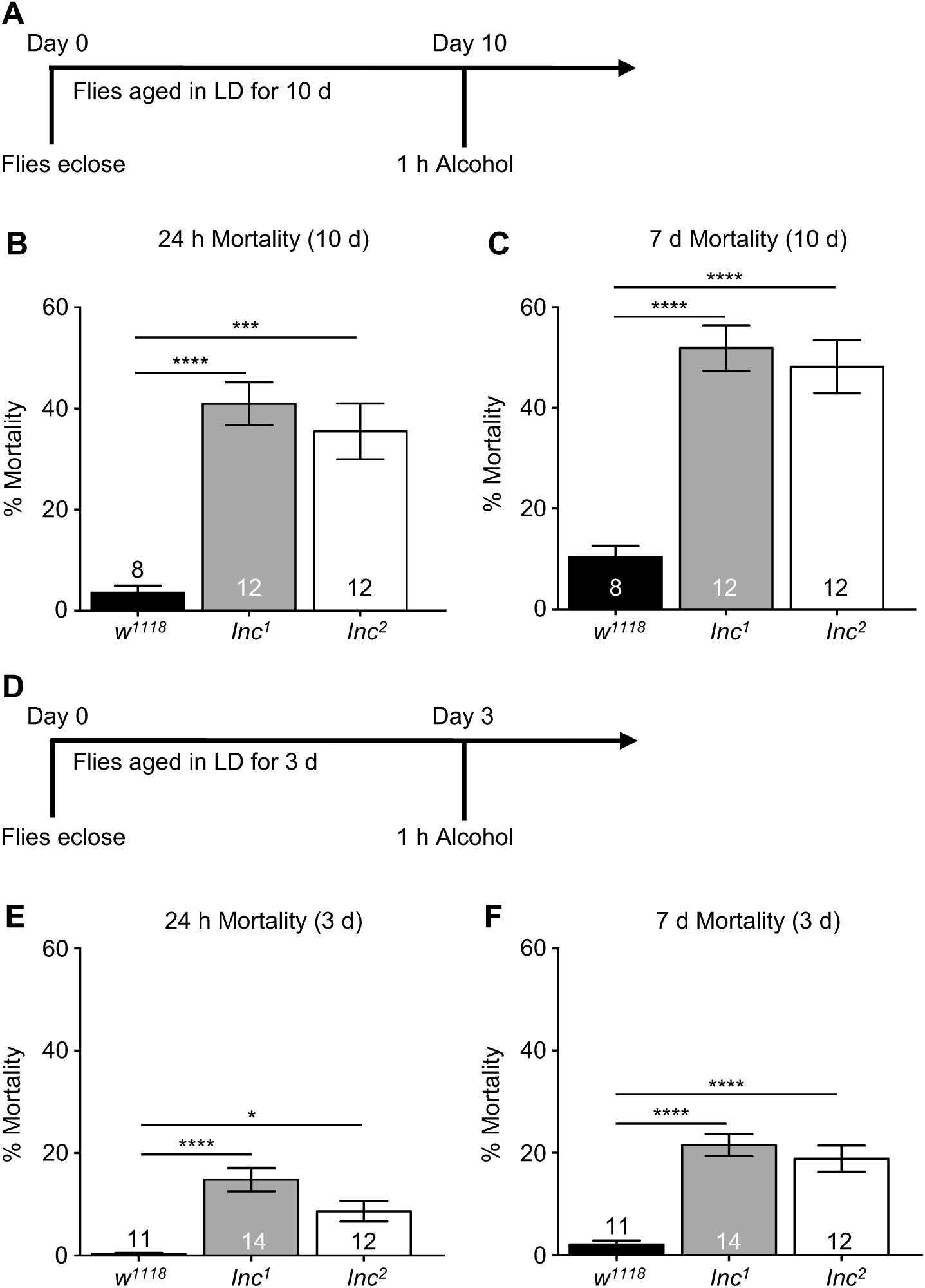
Short sleeping flies have significantly increased mortality following a single exposure to alcohol. A) The impact of sleep loss due to mutations in *inc^1^* and *inc^2^* on alcohol-induced mortality was assessed. 10 d *w^1118^*, *inc^1^* and *inc^2^* flies were exposed to 50% alcohol vapor for 1 h at ZT 9 with mortality assessed every 24 h for 7 d following exposure to alcohol. B) *inc^1^* and *inc^2^* mutant flies showed significantly increased mortality 24 h following exposure to alcohol compared to *w^1118^* control flies (ANOVA: *F_2,29_* = 16.46, *p* < 0.0001). C) Mortality in *inc^1^* and *inc^2^* flies continued to rise 7 d following the initial exposure to alcohol (ANOVA: *F_2,29_* = 20.87, *p* < 0.0001). D) To determine whether the alcohol-induced mortality in 10 d *inc^1^* and *inc^2^* flies was due to accumulation of sleep deficits, 3 d *inc^1^*, *inc^2^* and *w^1118^* control flies were exposed to 50% alcohol vapor for 1 h at ZT 9 with mortality assessed every 24 h for 7 d after alcohol exposure. E) The rate of mortality 24 h following exposure to alcohol was significantly higher in 3 d *inc^1^* and *inc^2^* flies compared to *w^1118^* controls but significantly lower than that of 10 d *inc^1^* and *inc^2^* flies (ANOVA: *F_2,34_* = 15.03, *p* < 0.0001). F) Compared to *w^1118^* controls, the mortality in *inc^1^* and *inc^2^* mutant flies continued to rise 7 d following the alcohol exposure but remained lower than that of 10d *inc^1^* and *inc^2^* mutant flies (ANOVA: *F_2,34_* = 25.01, *p* < 0.0001). These results support the hypothesis that sleep loss accumulated with age exacerbates susceptibility to alcohol-induced toxicity.

### Pharmacologically increasing sleep ameliorates alcohol-induced mortality in populations with sleep phenotypes

Previously we found that circadian arrhythmia and aging significantly increase alcohol-induced mortality^61^ mirroring human populations such as shift-workers and the elderly with sleep disturbances. If accrued sleep loss is the driving force for the observed alcohol-induced mortality in *inc* mutants, we hypothesized that increasing sleep in the *inc^1^* and *inc^2^* mutants should decrease mortality following exposure to alcohol. To pharmacologically increase sleep, *inc^1^* and *inc^2^* mutant flies were raised on standard *Drosophila* media for 9 d and then transferred to media containing the GABA_A_agonist THIP which has previously been shown to pharmacologically increase sleep in *Drosophila* ^77–79^. Following THIP exposure, *inc^1^* and *inc^2^* flies were given a single 1 h exposure to alcohol (Figure 6A). THIP exposure significantly reduced mortality 24 h and 7 d following alcohol exposure in both *inc^1^* and *inc^2^* mutant flies compared to non-THIP exposed *inc^1^* and *inc^2^* mutants (Figure 6B, ANOVA: *F_3,44_* = 13.27, *p* < 0.0001 and Figure 6C, ANOVA: *F_2,44_* = 12.83, *p* < 0.0001 respectively). However, THIP has dual effects as an analgesic and anxiolytic, and has been tested as a treatment for both alcohol use disorders as well as insomnia^80^. Potentially, as an agonist for GABA_A_receptors, THIP may be affecting alcohol-receptor interactions to affect mortality rather than through its pharmacological induction of sleep. To determine whether acute THIP interactions decreased alcohol-induced mortality by altering alcohol-receptor interactions rather than through increased sleep prior to alcohol exposure, we fed 10 d *inc^1^*, *inc^2^* and *w^1118^* flies 0.1 mg/mL THIP for 1 h at ZT 7-8 and then exposed them to 50% alcohol vapor. There were no differences in mortality between THIP-fed *inc^1^* and *inc^2^* flies and non-THIP fed flies following alcohol exposure (Figure 6E, ANOVA: *F_5,46_* = 50.88, *p* < 0.0001 and Figure 6F, ANOVA: *F_5,46_* = 55.83, *p* < 0.0001). These results are consistent with the hypothesis that increased sleep prior to binge-like alcohol exposure buffers the toxic effects of alcohol.

**Figure 6:**
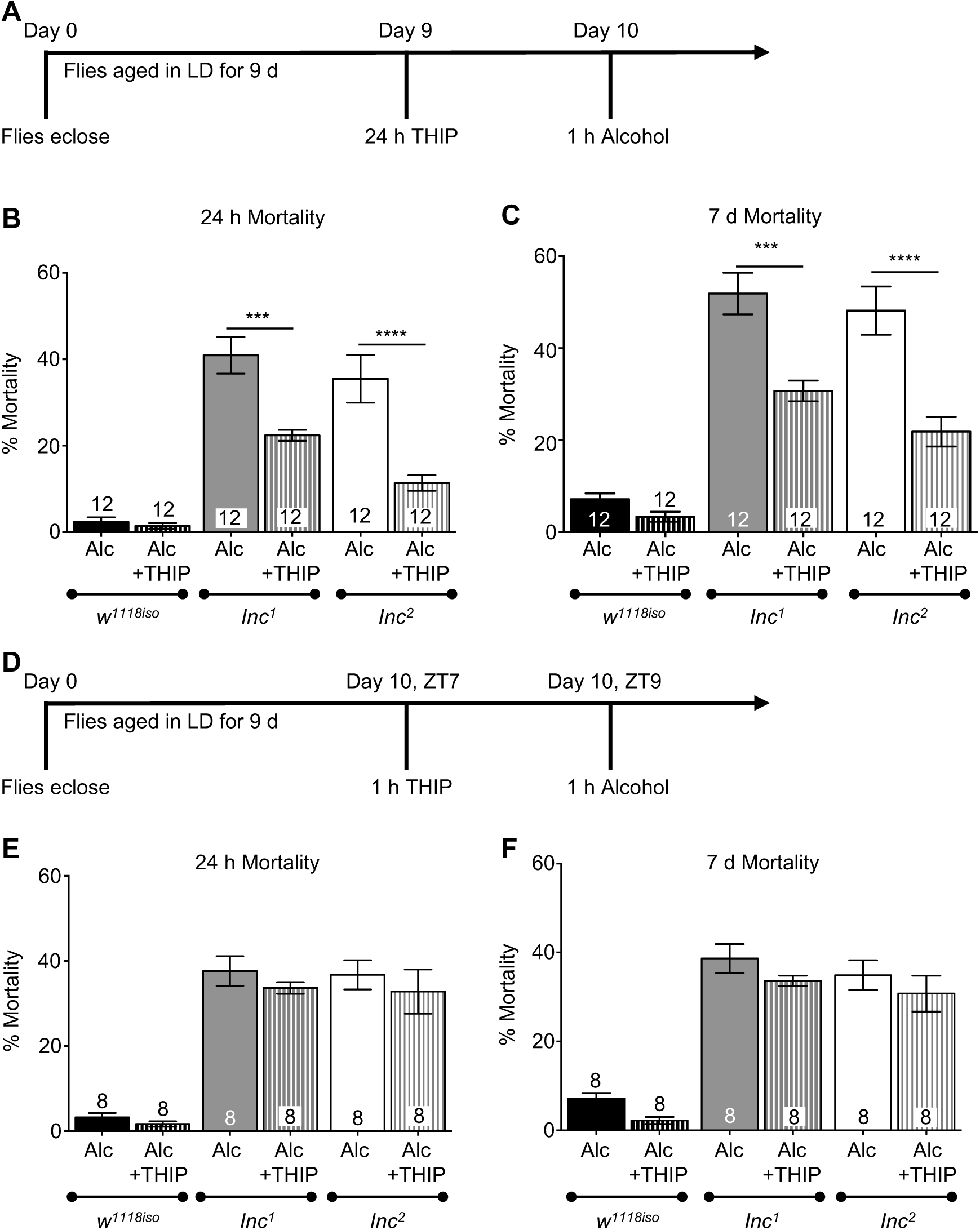
Pharmacologically increasing sleep in *insomniac* mutants ameliorates mortality following alcohol exposure. A) To determine whether increased sleep buffered against the toxic effects of alcohol, *inc^1^*, *inc^2^* and *w^1118^* flies were aged in LD cycles for 9 d and transferred to media containing 0.1 mg/mL THIP for 24 h. On day 10, flies were exposed to 1 h of 50% alcohol vapor at ZT 9 with mortality being assessed every 24 h for 7 d following exposure. B) Significantly reduced mortality was observed in THIP-fed *inc^1^* and *inc^2^* flies 24 h following alcohol exposure compared to *inc^1^* and *inc^2^* flies given alcohol alone (ANOVA: *F_3,44_* = 13.27, *p* < 0.0001). C) Although mortality rose in THIP-fed *inc^1^* and *inc^2^* flies 7 d following alcohol exposure, the percent of THIP-fed *inc^1^* and *inc^2^* flies was still significantly lower than non-THIP fed *inc^1^* and *inc^2^* flies at 7 d after exposure to alcohol (ANOVA: *F_2,44_* = 12.83, *p* < 0.0001). D) To determine whether the decreased mortality observed in THIP-fed flies was due to buildup of GABA-A receptor tolerance, *inc^1^*, *inc^2^* and *w^1118^* flies were aged in LD cycles for 10 d and transferred to media containing 0.1 mg/mL THIP for 1 h at ZT 7. At ZT 9, the flies were exposed to 1 h of 50% alcohol vapor with mortality being assessed every 24 h for 7 d following exposure. E) No significant differences in mortality were observed between THIP-fed *inc^1^* and *inc^2^* flies 24 h following alcohol exposure (ANOVA: *F_5,46_* = 50.88, *p* < 0.0001). F) No significant differences in mortality were observed in THIP-fed *inc^1^* and *inc^2^* flies compared to non-THIP fed flies 7 d following alcohol exposure (ANOVA: *F_5,46_* = 50.88, *p* < 0.0001).

### Pharmacologically increasing sleep, independent of circadian rhythmicity, decreases alcohol-induced mortality

In *Drosophila*, the circadian clock can be rendered non-functional using environmental manipulation by housing the flies in constant light. Constant light (LL) is sufficient to dampen molecular oscillations and abolish circadian rhythms in locomotor activity, memory formation and the rhythm in alcohol-induced loss-of-righting reflex ^51, 63, 81–85^. We have previously shown environmental disruption of circadian function exacerbates alcohol sensitivity and mortality^48, 61^. Along with a disrupted circadian clock, we found that 10 d CS flies in LL have significantly lower total sleep, specifically less sleep during the subjective night compared to 10 d CS flies in LD (Figure 7A; *t*_(236)_ = 4.46, *p* < 0.0001) and Figure 7B, ANOVA: *F_3,442_* = 27.31, *p* < 0.0001), consistent with mis-timed sleep due to circadian dysfunction. Flies housed in LL had significantly higher number of sleep bouts in both the subjective day and night (Figure 7C, ANOVA: *F_3,442_* = 98.99, *p* < 0.0001), although the sleep bout length was significantly shorter than flies housed in LD resulting in the decrease in total sleep (Figure 7D, ANOVA: *F_3,442_* = 78.45, *p* < 0.0001). As a first step to separate the effects of sleep from the effects of circadian disruption on alcohol toxicity, we characterized the effects of THIP on sleep for flies housed in LL. As expected, flies housed on THIP containing food in constant light slept significantly more than flies on regular *Drosophila* food in LL (Figure 7E – H; Mean sleep time per day: *t*_(233)_ = 29.43, *p* < 0.0001; Mean sleep time, day vs night, ANOVA: *F_3,463_* = 437.1, *p* < 0.0001; number of sleep bouts, ANOVA: *F_3,463_* = 358.1, *p* < 0.0001; Mean sleep bout length, ANOVA: *F_3,463_* = 277.6, *p* < 0.0001). To separate the role of sleep from circadian regulation in mediating alcohol toxicity following a repeat binge-like alcohol exposure, we increased sleep in flies in LL as they remained under conditions of circadian disruption. 10 d LL flies were maintained on medium containing THIP for 48 hours prior to exposure to the first of three exposures of 40% alcohol vapor (Figure 7I). LL flies housed on THIP containing food prior to alcohol exposure had a significantly lower mortality rate than those exposed to alcohol vapor alone (Figure 7J, ANOVA: *F_3,36_* = 132.6, *p* < 0.0001). However, LL flies given a short exposure to THIP followed by alcohol exposure exhibited no differences in mortality compared to LL flies exposed to alcohol alone (Figure 7K, ANOVA: *F_3,36_* = 92.26, *p* < 0.0001). These results suggest that increased sleep is sufficient to ameliorate mortality following repeated binge-like alcohol exposure even under conditions of circadian disruption.

**Figure 7:**
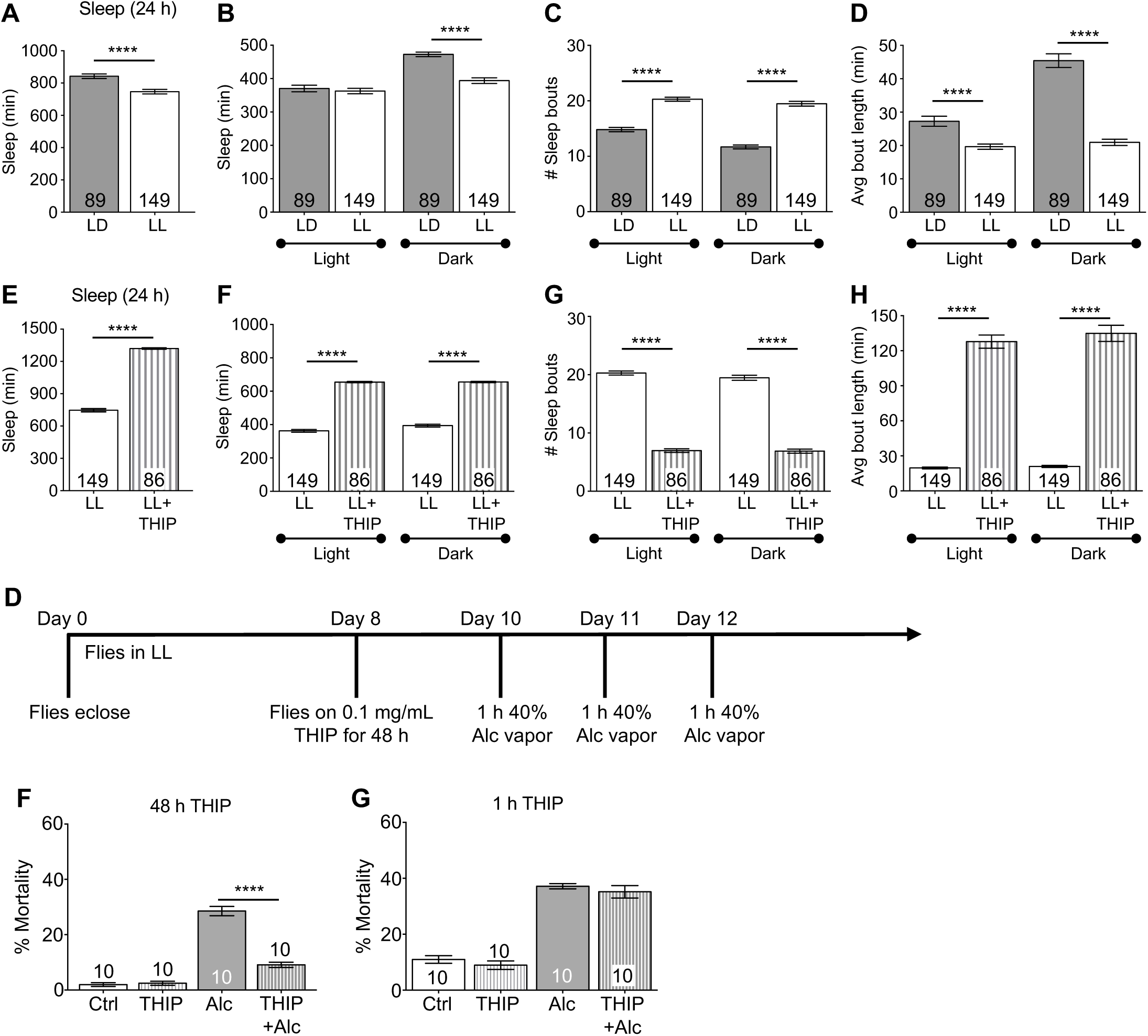
Pharmacologically increasing sleep in flies that are circadianly disrupted reduces mortality following repeated exposures to alcohol. A-D) Comparison of sleep profiles between 10 d WT CS flies housed in LD cycles and 10 d flies housed in constant light conditions (LL). Compared to control WT flies grown in a 12:12 h LD cycle, 10 d WT CS flies grown in LL exhibit significantly decreased average sleep time per day (A, [*t*_(236)_ = 4.46, *p* < 0.0001)]), decreased sleep during both the light and dark cycle (B, ANOVA: *F_3,442_* = 27.31, *p* < 0.0001), increased number of sleep bouts (C, ANOVA: *F_3,442_* = 98.99, *p* < 0.0001) and decreased bout duration (D, ANOVA: *F_3,442_* = 78.45, *p* < 0.0001). 10 d LL flies fed 0.1 mg/mL THIP exhibit significantly higher quiescence compared to 10 d LL flies on control media with significantly longer total sleep time per day (E, *t*_(233)_ = 29.43, *p* < 0.0001), increased sleep time during the light and dark cycles (F, ANOVA: *F_3,463_* = 437.1, *p* < 0.0001), decreased number of sleep bouts (G, ANOVA: *F_3,463_* = 358.1, *p* < 0.0001) and increased bout duration (H, ANOVA: *F_3,463_* = 277.6, *p* < 0.0001). I) To determine whether increasing sleep was sufficient to ameliorate alcohol-induced mortality under conditions of circadian disruption, WT CS flies are housed in LL upon eclosion and transferred to media containing 0.1 mg/mL THIP on day 8 for 48 h. On days 10, 11 and 12, flies were subjected to a three-exposure repeated binge-like alcohol paradigm with 1 h alcohol exposure (50% alcohol vapor) occurring at ZT 9 and mortality was assessed. J) Alcohol-induced mortality in THIP-fed LL flies was drastically reduced compared to control LL flies (ANOVA: *F_3,36_* = 132.6, *p* < 0.0001). K) Reduced mortality following THIP exposure is not due to increased tolerance as 1 h exposure to 0.1 mg/mL immediately prior to alcohol exposure had no significant effect on reducing alcohol-induced mortality (ANOVA: *F_3,36_* = 92.26, *p* < 0.000).

### Increasing sleep buffers age-related susceptibility to alcohol-induced mortality

Aging is accompanied by the breakdown of circadian rhythmicity at the cellular, metabolic and physiological levels as well as disruptions in sleep architecture^61, 86–90^. In recent years, chronic and binge alcohol consumption in middle-aged and older adults has significantly increased^91, 92^ with more than 75% of the alcohol-induced poisoning deaths occurring in these age groups^11, 14^. More than 10% of older adults engage in binge drinking behavior^91^. Given that the aging population is expected to double by 2050^93^, it is necessary to identify ways to treat or ameliorate alcohol toxicity in middle-aged and older individuals. In previous studies, we have shown that aging exacerbates alcohol sensitivity and mortality^61^. Middle-aged flies (20 d) exhibit shorter sleep times compared to younger flies (^40^; Figure 8A, (*t*_(147)_ = 6.54, *p* < 0.0001) and Figure 8B, ANOVA: *F_3,294_* = 38.12, *p* < 0.0001). While we found no differences in total sleep amount during the night between 10 and 20 d flies, 20 d flies had a significantly greater number of sleep bouts with shorter duration reflecting decreases in sleep consolidation (Figure 8C, ANOVA: *F_3,294_* = 19.21, *p* < 0.0001 and Figure 8D; ANOVA: *F_3,294_* = 38.12, *p* < 0.0001). We tested whether pharmacologically increasing sleep in middle-age was sufficient to overcome the age-related increase in mortality following repeated binge-like exposures to alcohol. We pharmacologically induced sleep in 20 d CS flies by housing them on 0.1 mg/mL THIP for 24 h after which they were given a 1 h alcohol exposure for 3 consecutive days (Figure 8I). 20 d THIP-fed flies slept significantly more than control 20 d flies (Figure 8E – 8H; Avg total sleep/day: *t*_(89)_ = 13.05, *p* < 0.0001; Avg sleep time, day vs night, ANOVA: *F_3,178_* = 91.72, *p* < 0.0001; number of sleep bouts, ANOVA: *F_3,178_* = 60.24, *p* < 0.0001; Sleep bout length, ANOVA: *F_3,178_* = 113.0, *p* < 0.0001). Middle-aged flies housed on THIP containing food prior to the repeated binge-like alcohol exposures had significantly lower rates of mortality than those exposed to alcohol alone (Figure 8J, ANOVA: *F_3,44_* = 243.8, *p* < 0.0001). The decreased mortality observed in THIP-fed flies was not due to increased alcohol tolerance from THIP interactions as 20 d flies given THIP for 1 h at ZT 7 followed by alcohol exposure at ZT 9 show mortality rates similar to 20 d flies exposed to alcohol alone (Figure 8K, ANOVA: *F_3,30_* = 23.09, *p* < 0.0001). These results suggest that increased sleep is sufficient to ameliorate mortality following repeated alcohol exposures in middle-aged flies that have both circadian and sleep disruption.

**Figure 8:**
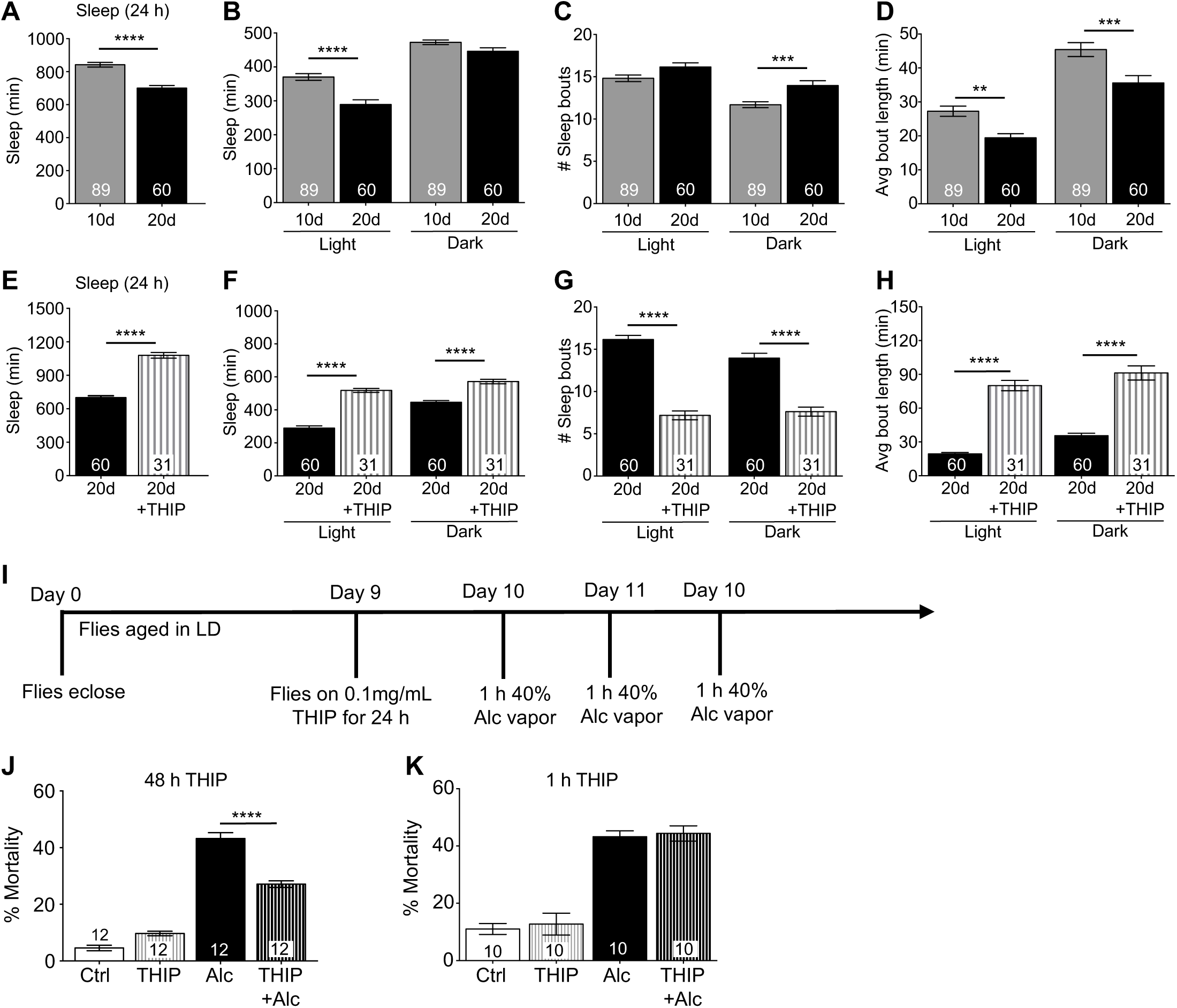
Pharmacologically increasing sleep in aging ameliorates alcohol-induced mortality. Compared to young 10 d flies, 20 d flies grown in LD cycles exhibit significantly decreased average sleep time per day (A), decreased sleep during both the light and dark cycle (B), increased number of sleep bouts (C) and decreased bout duration (D). 20 d flies fed 0.1 mg/mL THIP exhibit significantly higher quiescence compared to 20 d flies on control media with significantly longer total sleep time per day (E) increased daytime and nighttime sleep. I) Flies are grown in 12 h light: 12 h dark cycles and transferred to media containing 0.1 mg/mL THIP on day 19 for 24 h. On days 20, 21 and 22, flies were subjected to a 3-exposure repeated binge-like alcohol paradigm with 1 h alcohol exposure (40% alcohol vapor) occurring at ZT 9 and mortality was assessed. J) Increased quiescence in 20 d THIP-fed flies significantly ameliorated alcohol-induced mortality compared to 20 d flies fed control media (ANOVA *F*_3,44_ = 243.8, *p* < 0.0001). K) 20 d flies exposed for 1 h to 0.1 mg/mL THIP immediately prior to alcohol exposure had no significant effect on reducing alcohol-induced mortality [*t*(9) = 1.047, *p* = 0.3223].

### Sleep deprivation inhibits long-term but not short-term tolerance

*Drosophila* exhibit drug tolerance with repeat alcohol exposures in which the behavioral response to subsequent exposures of alcohol is lessened similar to that observed in rodent models and humans. At the behavioral level, functional tolerance results in a decreased sensitivity to alcohol during subsequent exposures with increased alcohol concentrations or longer alcohol exposures necessary to induce sedation^33, 94, 95^. In flies, rapid tolerance develops after a single alcohol exposure and can be observed during a second alcohol exposure 4 h or 24 h later^50, 96^. The development of functional alcohol tolerance is dependent upon changes in neural plasticity rather than changes in the metabolism or clearance of alcohol^50, 53–55, 96, 97^. Changes in neural plasticity associated with drug and alcohol tolerance share features in common with synaptic plasticity observed in learning and memory^98–100^. Potentially, sleep loss affects the development of alcohol tolerance as sleep deprivation disturbs memory formation as seen across invertebrate and vertebrate species^101–103^. To investigate the effect of sleep deprivation on tolerance formed after a single alcohol exposure, 10 d wild-type flies were sleep deprived for 24 hours (ZT 3.5 – ZT 3.5) and given a pre-exposure of 50% alcohol vapor for 30 minutes (ZT 4.5; Figure 9A). Pre-exposed sleep deprived flies and sleep deprived naïve flies were exposed to alcohol 4 h later at ZT 9 during which sedation was measured (Figure 9A). Similarly, non-sleep deprived flies were pre-exposed to alcohol with responses compared during a second alcohol exposure to naïve flies. Non-sleep deprived flies demonstrated robust 4 h alcohol tolerance with significant increases observed in the time necessary for 50% of the flies to reach sedation compared to naïve flies (Figure 9B, ANOVA: *F_3,20_* = 49.62, *p* < 0.0001 and Figure 9C). Surprisingly, sleep deprived flies also demonstrated robust 4 h alcohol tolerance (Figure 9B and C) suggesting that sleep disruption does not affect the cellular signaling mechanisms necessary for the formation of 4 h tolerance.

**Figure 9:**
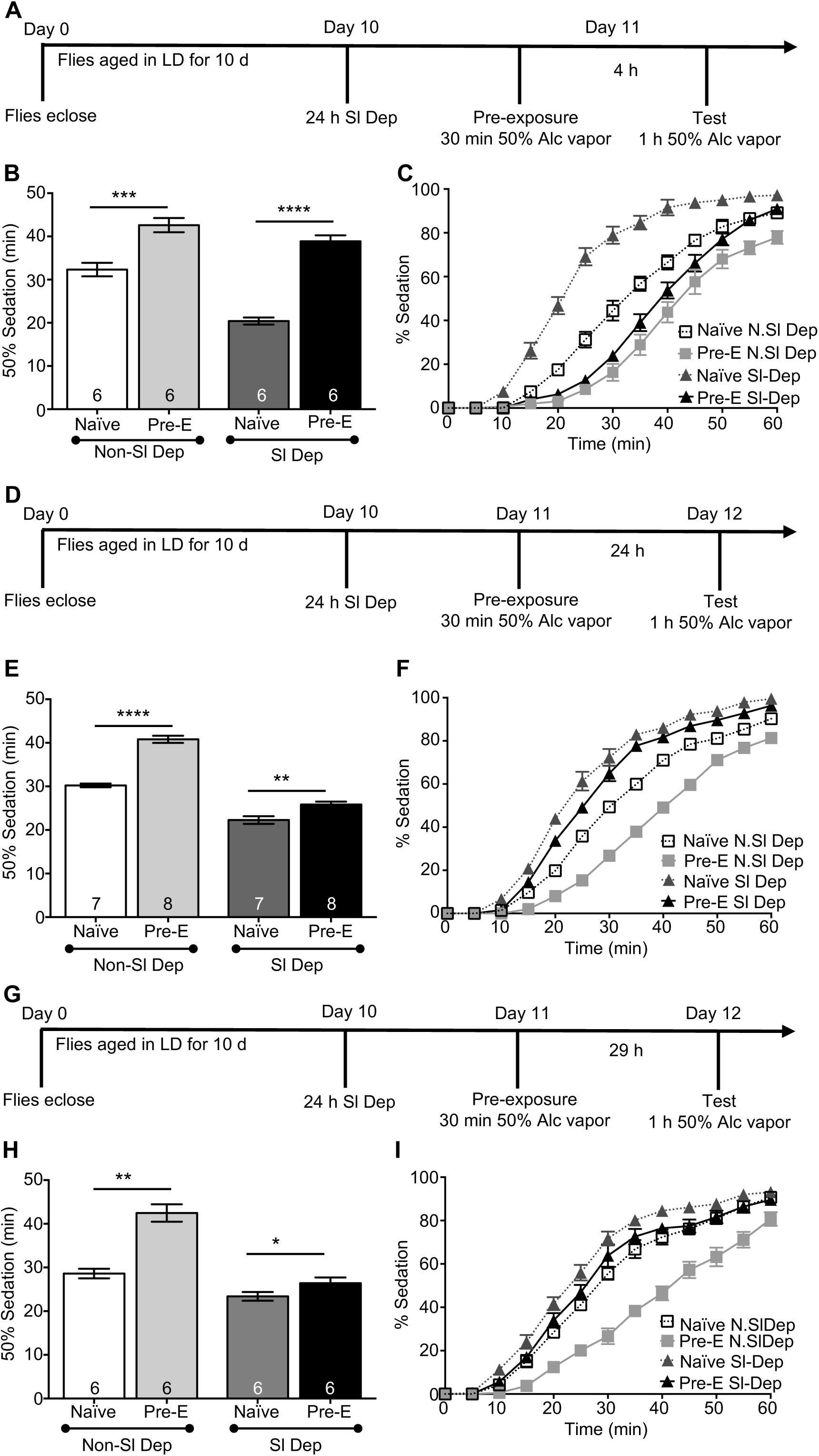
Sleep deprivation differentially affects short-term and long-term functional tolerance. **A-C)** Effect of 24 h sleep deprivation on short-term functional alcohol tolerance. A) WT CS flies were aged in 12:12 h LD cycle and sleep deprived for 24 h on day 10. On day 11, flies were exposed to 50% alcohol vapor for 30 min and tested 4 h later by exposing to 50% alcohol vapor for 1 h with sedation being measured. B) 24 h sleep deprivation has no significant effect on the development of short-term acute alcohol tolerance. Both sleep deprived and non-sleep deprived flies exhibited alcohol tolerance (ANOVA *F*_3,20_ = 49.62, *p* < 0.0001). C) Complete time course of alcohol exposure showing percent of flies exhibiting sedation for 10 d old sleep-deprived and non-sleep deprived flies. D-F) Effect of sleep deprivation on development of long-term functional alcohol tolerance. WT CS flies were aged in 12:12 h LD cycle and sleep deprived for 24 h on day 10. On day 11, flies were exposed to 50% alcohol vapor for 30 min and tested 24 h later by exposing to 50% alcohol vapor for 1 h at ZT 9 with sedation being measured. E) Both sleep-deprived and non-sleep deprived flies exhibited alcohol tolerance (ANOVA *F*_3,26_ = 125.7, *p* < 0.0001) with 24 h sleep deprivation significantly dampens development of long-term functional alcohol tolerance. F) Complete time course of alcohol exposure showing percent of flies exhibiting sedation for 10 d old sleep-deprived and non-sleep deprived flies.

To determine the effect of sleep deprivation on the formation of long-term alcohol tolerance, flies were sleep deprived for 24 hours (ZT 7.5 – ZT 7.5) and given a pre-exposure of 50% alcohol vapor for 30 minutes (ZT 8.5) and tested 24 hours later at ZT 9 (Figure 9D). Groups of non-sleep deprived flies were handled concurrently. When flies were tested 24 h after the initial alcohol exposure, sleep-deprived flies demonstrated significantly less tolerance to alcohol with the time to sedation similar to sleep-deprived naïve flies while non-sleep deprived flies demonstrated robust long-term tolerance with response times significantly different than naïve flies (Figure 9E, ANOVA: *F_3,26_* = 125.7, *p* < 0.0001 and Figure 9F). Although our previous research found that tolerance was not modulated by the circadian clock, we verified the effect on long-term tolerance by exposing flies to alcohol at the same time that we observed the formation of 4 h tolerance in sleep-deprived flies (Figure 9G). Sleep-deprived flies pre-exposed to alcohol at ZT 8.5 and then subsequently exposed to alcohol at ZT 9 the following day also exhibited little or no alcohol tolerance, while non-sleep deprived flies exhibited significant long-term tolerance (Figure 9H, ANOVA: *F_3,20_* = 0.92, *p* = 0.4488 and Figure 9I). Thus, acute sleep deprivation prior to alcohol exposure inhibits the expression of alcohol tolerance 24 h following the initial alcohol pre-exposure while no effect is observed on the development of short-term tolerance expressed 4 h after the initial exposure. These results are consistent with the hypothesis that different molecular mechanisms underlie the development of short-term and long-term rapid alcohol tolerance similar to the differences in formation of short and long-term memory.

## Discussion

Research from our lab and others suggests a bidirectional relationship between clock dysfunction and the onset and severity of alcohol-related pathologies^18, 48, 51, 104, 105^. Social jet lag, large shifts in sleep timing between the weekday and the weekend is observed in numerous populations, including individuals on shift and rotating schedules^106, 107^ and is strongly correlated with increased alcohol use^108, 109^. Due to the long working hours, rotating schedules and work-associated stress, many individuals report using alcohol as a sleep aid^64, 65, 110, 111^ which can eventually lead to an increased number of binge drinking episodes and other detrimental effects associated with alcohol abuse^64, 65, 67, 112^. Previous studies from our lab found that the circadian clock modulates alcohol sensitivity and toxicity and that circadian dysfunction significantly increases the behavioral sensitivity to alcohol and mortality following acute and repeated alcohol exposures^48, 51^. In humans, differences in individual chronotype also appears to modulate alcohol use and its associated pathologies. Individuals expressing an “evening chronotype” report significantly increased alcohol use^113–124 125^. Interestingly, individuals with an evening chronotype also have lower quality of sleep and increased greater daytime fatigue^126, 127^. However, it has been difficult to detangle the effects of circadian dysfunction from the effects of altered sleep on alcohol use.

Sleep disorders and sleep disturbances have become increasingly prevalent in modern society with longer working hours, irregular work schedules and the prevalence of electronics, affecting more than 35% of adults and 70% of teenagers^27, 74, 128–131^. Insufficient sleep exacerbates the risk of developing chronic diseases and health problems including cancer, diabetes, neurodegenerative and psychiatric disorders^132–136^. Consequently, we investigated the effects of sleep loss on the alcohol sensitivity and toxicity using *Drosophila melanogaster* to dissect the interactions between sleep deprivation and alcohol sensitivity and mortality.

We found that acute (24 h) sleep deprivation significantly increased sensitivity and mortality in young flies following a single binge-like exposure to alcohol. Most of the observed increase in mortality following alcohol exposure occurred within 24 h following alcohol exposure. These effects were independent of stress or injury as 48 h recovery sleep prior to alcohol minimized alcohol-induced mortality. The increases in sensitivity and mortality were also independent of changes in metabolic tolerance as there were no differences between sleep deprived and non-sleep deprived flies in alcohol absorbance or clearance. Thus, sleep deprivation changes both immediate alcohol sensitivity and acute alcohol toxicity after a single binge-like alcohol exposure. Our data underlines the phylogenetic conservation across species showing a correlation between sleep loss and alcohol behaviors. Studies from rodents and humans outline a correlation between sleep loss and increased severity of alcohol behavioral responses including increased alcohol intake, accelerated development of alcohol abuse, dependence and relapse following alcohol abstinence^137–140^. In mice, alcohol dose dependently increases hyperactive locomotor activity in open-field tests with acute sleep deprivation for 48 h abolishing these stimulatory effects^141^. Insufficient sleep (< 8 h per night) is correlated with increased number of drinking sessions in adolescents and young adults^142–144^. College-aged students are considered a vulnerable population for risk-taking behaviors and multiple studies show a strong correlation between poor sleep quality and excessive alcohol intake and the accompanying consequences for mental health and academic performance including increased rates of depression, anxiety, psychological stress and academic issues in these students^66, 145, 146^. Insufficient and poor quality sleep also appear to predict the onset of alcohol abuse and its adverse consequences^68, 147–150^. Sleep disturbances observed in children 3-5 years of age predicted the early onset of alcohol use at ages 12-14^151^. This is particularly harmful because recovering alcoholics who used alcohol as a sleep aid are three times more likely to relapse in 12 months^22, 23, 152^. Altogether, these studies emphasize disturbed sleep as a potent risk factor for the initiation of alcohol use, escalation of problems associated with alcohol abuse and hindrance of recovery from alcohol-use disorders. With the genetic tools and mutants available, *Drosophila* provides a suitable model system to test the relationship between chronic sleep disturbances and alcohol induced pathologies. Using flies with mutations in the *insomniac* gene (*inc*) that provide a model mirroring chronic sleep restriction, we found that *inc* mutants have significantly increased mortality following alcohol exposure than background controls. Moreover, we found that alcohol exposure is more lethal in 10 d old *inc*-mutant flies compared to 3 d old flies, although wild type flies in either age groups show little alcohol-induced mortality. *Inc* mutant flies were surprisingly less sensitive to the sedative effects of alcohol compared to their background controls supporting previous research that the different physiological consequences of alcohol can be regulated separately. Although, the mechanism through which sleep buffers alcohol toxicity is unknown, it is possible that the changes in oxidative stress in the *inc* mutant flies may contribute to the change in alcohol toxicity. The *inc* gene seems necessary for mediating the oxidative stress response as reducing *inc* both globally and neuronally significantly increases mortality following a single injection to paraquat, a common inducer of oxidative stress^76, 153^. Support for this hypothesis is found in previous research demonstrating that pharmacologically increasing sleep in *inc*-mutant flies using gaboxadol significantly decreased the sensitivity to paraquat-induced oxidative stress^153^. Sleep loss has also been shown to increase reactive oxygen species in the gut^154^ raising the possibility that peripheral mechanisms also contribute to increased alcohol toxicity. As changes in sleep potentially impact multiple physiological processes in the central nervous system as well as in peripheral organs, the precise mechanism(s) through which sleep buffers alcohol toxicity will undoubtedly be the focus of future studies.

Pharmacologically increasing sleep alone in circadianly disrupted and middle-aged wild-type flies was sufficient to significantly reduce alcohol-induced mortality. Gaboxadol increased total sleep duration as well as significantly increasing sleep bout length suggesting a greater consolidation of sleep. Both increased total sleep and increased sleep consolidation suggest that improved sleep quality could aid in mitigating alcohol-induced pathologies. Although there have been few studies examining the relationship between sleep health and alcohol toxicity, sleep loss or decreased sleep consolidation has been shown to reduce reproductive output, accelerate aging and increase the accumulation of reactive oxygen species and death in flies^154, 155^. In humans, increasing sleep in adolescents is correlated with decreased risk of emotional and cognitive disruption as well as lowered risk of obesity^156^. Also, increasing sleep by 30 minutes for 3 days over the weekend in healthy industrial workers and individuals susceptible to obesity significantly increased insulin sensitivity and had a restorative effect of sleep on metabolic homeostasis^157, 158^. Finally, increasing sleep in older adults significantly improves performance on visual tasks and stabilizes memory recall^159^. Although more specific research needs to be done assessing the direct effects of increased sleep on alcohol toxicity in vulnerable groups, these data suggest a role for sleep as a buffer to protect against the toxic effects of alcohol in populations vulnerable to chronic sleep loss as aged adults and shift workers.

The development of acute tolerance to alcohol is a distinct and critical behavioral metric used to gauge the propensity for alcohol dependence and abuse^160^, that can be separated from alcohol sensitivity and alcohol toxicity. Similar to mammals, an acute exposure to a high concentration of alcohol induces functional tolerance in *Drosophila* at the behavioral^50, 96, 161^ and the molecular levels^97, 162–165^. Functional alcohol tolerance is dependent on changes in neuronal strength and connectivity or synaptic plasticity^97, 162, 163^. Consistent with previous findings, we observed tolerance 4 hours and 24 hours following a short pre-exposure to alcohol vapor^50, 51^. We found that sleep deprivation abolished the development of tolerance at 24 hours but had no effect on tolerance at 4 hours. Potentially, acute sleep deprivation selectively impairs the cellular and molecular processes necessary for encoding long-term rapid tolerance to alcohol without severe disruption of those mechanisms necessary for the development of 4 hour tolerance. In fact, previous studies demonstrate altered expression of rapid tolerance in flies with mutations in genes necessary for learning and memory^46, 96, 166^. For example, the gene *dunce* (*dnc*) encodes a phosophodiesterase required for cAMP degradation and is necessary for behavioral and synaptic plasticity^167, 168^. Originally identified as a learning mutant^169, 170^, *dnc*-mutant flies exhibit significant sleep deficits^171^ and are incapable of forming rapid tolerance^172, 173^. Time-dependent differences in the effects of sleep deprivation can also be seen for memory with acute sleep deprivation affecting the consolidation of long-term but not short-term hippocampal dependent memory in mice^174, 175^. Together with support from existing research, the results from our studies suggest that sleep deprivation selectively impacts processes underlying synaptic plasticity to affect the development of long-term rapid tolerance. In conclusion, the results from our study start to dissociate the role of sleep in modulating alcohol toxicity from the regulation of alcohol neurobiology by the circadian clock. These results lay the groundwork for future studies and treatments considering sleep quality and sleep duration as an important component of alcohol use disorder and alcohol-induced pathologies.

## Disclosure Statement

Financial Disclosure: None

Non-financial Disclosure: None

An early version of this manuscript (the authors’ original version) prior to peer review may be found in bioRxiv.

## Acknowledgements

This work was supported by the National Institutes of Health, National Institute on Alcohol Abuse and Alcoholism grant R21AA021233.

